# Simulations of proposed mechanisms of FtsZ-driven cell constriction

**DOI:** 10.1101/737189

**Authors:** Lam T. Nguyen, Catherine M. Oikonomou, Grant J. Jensen

## Abstract

To divide, bacteria must constrict their membranes against significant force from turgor pressure. A tubulin homolog, FtsZ, is thought to drive constriction, but how FtsZ filaments might generate constrictive force in the absence of motor proteins is not well understood. There are two predominant models in the field. In one, filaments overlap to form complete rings around the circumference of the cell; as filaments slide against each other to maximize lateral contact, the rings constrict. In the other, filaments exert force on the membrane by a GTP-hydrolysis-induced switch in conformation from straight to bent. Here we developed software, ZCONSTRICT, for quantitative 3D simulations of Gram-negative bacterial cell division to test these two models and identify critical conditions required for them to work. We find that the avidity of lateral interactions quickly halts the sliding of filaments, so a mechanism such as depolymerization or treadmilling is required to sustain constriction by filament sliding. For filament bending, we find that a mechanism such as the presence of a rigid linker is required to constrain bending within the division plane and maintain the distance observed in vivo between the filaments and the membrane. We also explored the recent observation of constriction associated with a single FtsZ filament and found that it can be explained by the filament bending model if there is a rigid connection between the filament and the cell wall. Together, our work sheds light on the physical principles underlying bacterial cell division and informs future experiments to elucidate the mechanism of FtsZ.

## INTRODUCTION

To divide, bacterial cells must constrict their cytoplasmic membrane against the large opposing force of internal turgor pressure, but how this is done is not well understood. Rod-shaped bacteria surround their cytoplasmic membrane with a semi-rigid cell wall composed of long glycan strands running around the cell’s circumference, crosslinked by short peptides into a mesh-like network that provides the cell’s shape template (*1*–*3*). Gram-positive bacteria expand their thick wall into a septum across the cell during division, suggesting that the inward growth of cell wall pushing on the membrane from outside might be the main force generator. In Gram-negative bacteria by contrast, we showed recently that a thin cell wall is maintained throughout division (*4*). Simulating division of this Gram-negative system, we found that cell wall growth alone cannot cause constriction, even with a make-before-break mechanism in which complete hoops of new cell wall are built underneath the existing wall before old peptide bonds are cleaved. Rather, a constrictive force is required to initially relax the pressure on the existing wall, allowing incorporation of new cell wall hoops with smaller radius (*4*). How the cell might generate such a constrictive force is still unclear.

In many eukaryotic cells, a ring formed by actin filaments and myosin motor proteins is responsible for generating a constrictive force to divide cells (*5*), but motor proteins are absent from bacterial cells. Instead, the tubulin homolog FtsZ (*6*) has been proposed to generate a constrictive force at the midcell during division (*7, 8*), but how this might work remains debated. FtsZ forms filaments that align to the circumference of the cell (*9, 10*). The distance between the filaments and the membrane is 16 nm (*9, 10*), but what maintains this distance remains unclear. FtsZ filaments are connected to the membrane via FtsA and/or ZipA proteins that bind the C-terminus of FtsZ via a flexible linker (*11*–*13*). Besides its proposed role as a force generator, FtsZ is also thought to serve as a scaffold for the cell wall synthesis machinery (*7, 8*). Recently, FtsZ filaments were shown to treadmill around the cell circumference, and it has been suggested that such movement may help distribute new cell wall uniformly around the cell (*14, 15*).

There are two predominant conceptual models in the field regarding how FtsZ might generate a constrictive force: filament sliding and filament bending. Observation of bundles of FtsZ filaments *in vitro* (*16*–*25*) led to the hypothesis that FtsZ filaments overlap via lateral bonds to form a complete ring which then tightens to constrict the membrane. This was supported by Monte Carlo simulations that showed that lateral attractive interactions between filaments in a closed ring can lead to sliding-induced filament condensation and ring constriction (*26*). Later, an electron cryotomography (cryo-ET) study showed that complete rings of overlapping FtsZ filaments exist in dividing *Caulobacter crescentus* and *Escherichia coli* cells (*10*), providing experimental support for this filament sliding model. Recently, however, cryo-ET studies from our lab showed that in several species, initial constriction can occur when only a single FtsZ filament is visible, suggesting that FtsZ can generate a constrictive force without forming a complete ring (*27*). Alternatively, as FtsZ forms both straight and bent filaments *in vitro* (*28*–*34*), it has been proposed that filaments with a large bending stiffness can exert adequate force to constrict the membrane. Several lines of evidence showed that reconstituted FtsZ can deform liposomes in a process that depends on GTP hydrolysis (*35*–*38*). Consistent with this hypothesis, cryo-ET imaging in our lab showed that in dividing *C. crescentus* cells, FtsZ forms both straight and bent filaments. This result suggested an iterative pinching model in which FtsZ polymerizes into straight filaments which then hydrolyze GTP to bend, pinching the membrane, before depolymerizing to start another cycle (*9*).

To explore how FtsZ filaments might generate a constrictive force in the absence of a motor protein, here we developed software, ZCONSTRICT, to simulate the two predominant models, namely filament sliding and filament bending. For each, we identified conditions required for the model to work. For filament sliding, we found that: (1) a long-range attractive force between filaments is required to increase lateral contact; (2) since increasing lateral contact also increases avidity, mechanisms such as filament depolymerization or treadmilling are required to break avidity; and (3) overlapping filaments have to form a complete ring for sliding to generate ring tension. Exploring the filament bending model we found that: (1) a mechanism that causes bending such as GTP hydrolysis is required; (2) bending must be confined within the division plane, for instance by rigid linkers that prevent filament rolling; and (3) incomplete rings of filaments must be connected to the cell wall in order for the bending force to overcome the effect of turgor pressure. These results are summarized in Movie S1 (https://youtu.be/rw88IzRAxEM).

## RESULTS

We built a coarse-grained model of the system (Fig. 1) in which the membrane was modeled as a sheet of beads, originally forming a cylinder with a radius of 250 nm, consistent with the size of many Gram-negative bacterial cells. FtsZ filaments were initiated in a ring-like arrangement, with a ring radius of 234 nm, separating the filaments 16 nm away from the membrane, a distance that was observed experimentally (*9, 10*). The average length of the filaments was chosen to be 176 nm (40 monomers). For the filament sliding model, each FtsZ filament was modeled as a chain of beads connected by springs, with one bead representing one FtsZ monomer, and the filament was originally aligned to the membrane circumference. For the filament bending model, each filament was modeled as a chain of cubes, with each cube representing one FtsZ monomer. To model connections between the filament and the membrane, linkers were composed of two identical springs joined by a bead representing a linking protein, such as FtsA or ZipA. To reflect the flexibility of these linkers, the two springs were allowed to freely rotate around the joining bead without an energy cost. The cell wall was modeled as a grid of beads, originally having a radius of 265 nm, which reduced as the membrane constricted (see **Methods** for details).

**Figure 1:**
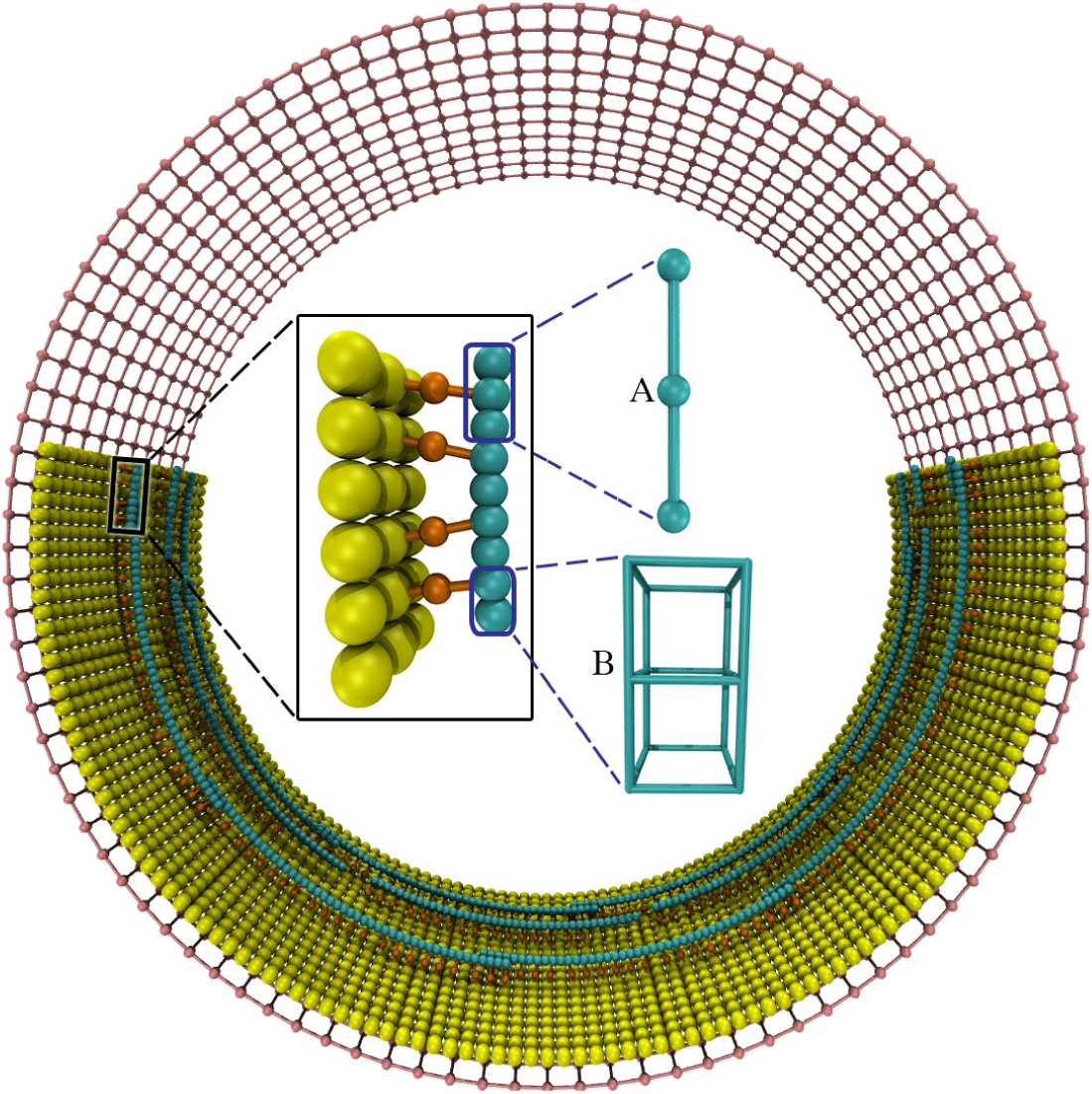
The coarse-grained model. The membrane (yellow) was modeled as a single layer of beads, originally forming a cylinder. Each FtsZ filament (cyan) was modeled as a chain of beads in the filament sliding model (A) or as a chain of cubes in the filament bending model (B). The filament was connected to the membrane via a set of linkers (orange), each composed of two springs (shown as rods) joined at a bead that represents an FtsA or a ZipA protein. The cell wall (pink) was modeled as a grid of beads, originally forming a cylinder surrounding the membrane. Note that the same colors are used for all following figures unless stated otherwise.

### Filament sliding model

#### Long-range interaction between filaments

We reasoned that for filament sliding to generate a constrictive force on the membrane, at least two conditions must be present. First, to induce sliding of filaments in lateral contact, a long-range attractive force has to exist between the filaments in order to increase the number of lateral bonds. Second, overlapping filaments have to form a complete ring in order for filament sliding to generate ring tension. (Note that we later verified this condition by observing that a broken ring failed to constrict the membrane.) We then constructed filaments that overlapped to form complete rings, separating adjacent rings by a distance of 20 nm (Fig. 2A). We then implemented a long-range Lennard-Jones potential between the beads on different filaments (see **Methods/Filament sliding model** for details). Simulations of this system showed that the rings failed to constrict the membrane but rather collapsed into a multi-layered bundle, with the high avidity between the filaments preventing sliding (Fig. 2B–D).

**Figure 2:**
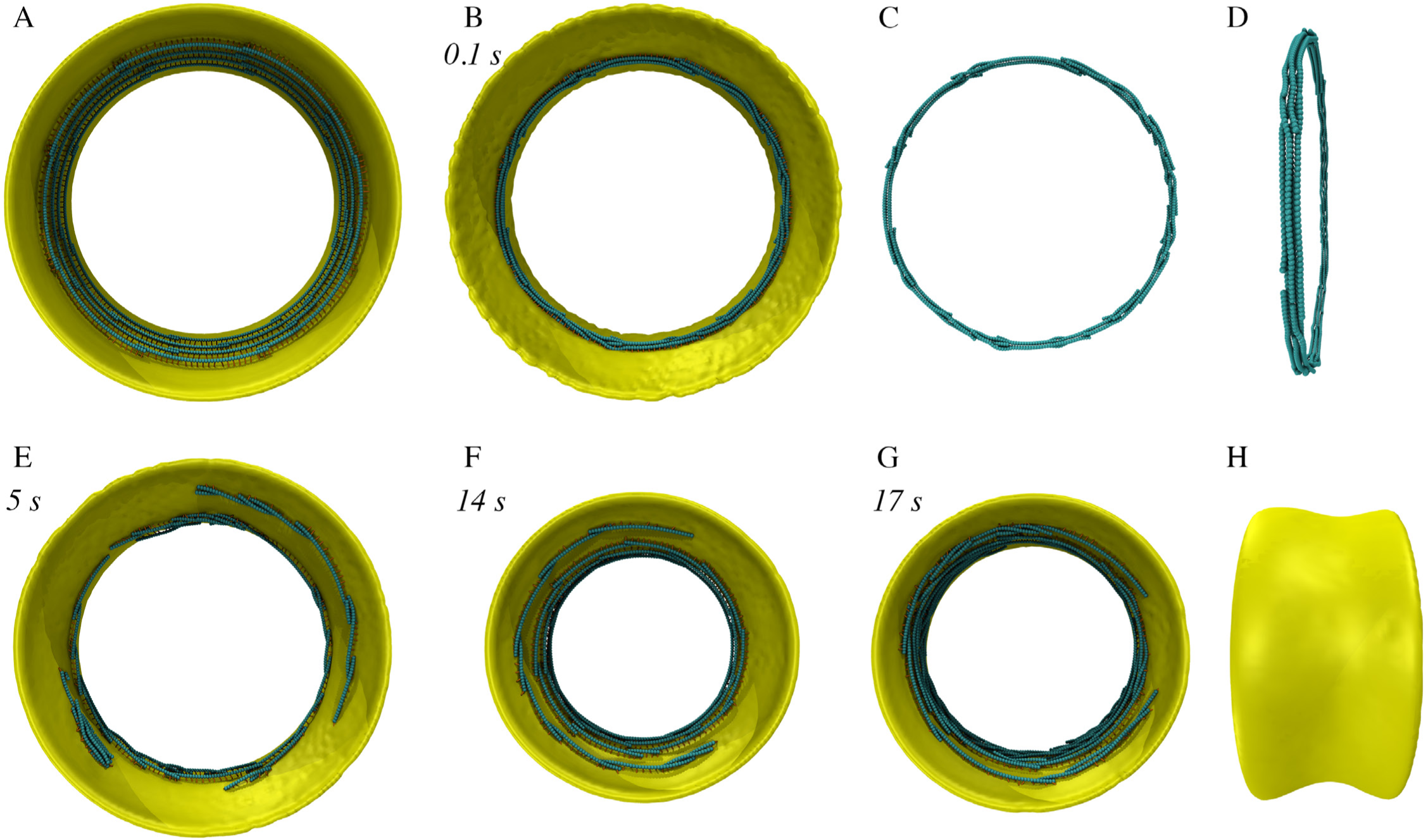
Simulation results of the filament sliding model. Italic font indicates simulation times. (A) Axial view of the initial system in which the FtsZ filaments overlapped to form complete rings. (B) Implementing a Lennard-Jones potential interaction between the filaments resulted in formation of a bundle of rings. (C) and (D) show axial and side views, respectively, of the ring bundle in (B). (E) Implementing depolymerization with a rate above the critical rate quickly resulted in loss of ring integrity. (F) With a depolymerization rate slower than *r*_*c*_, deep constriction occurred and rings broke much more slowly. (G) Removing depolymerization and implementing treadmilling also resulted in a deep constriction. (H) Side view of (G).

#### Ring separation

Since cryo-ET studies of dividing cells showed no evidence for multi-layered bundling (*9, 10, 27*), we judged this unlikely to occur in real cells, perhaps due to intervening proteins. Not knowing the *in vivo* mechanism, since filaments were originally arranged in separate rings, we simply calculated the Lennard-Jones potential only between filaments in the same ring. As a result, bundling was prevented. However, filament sliding only occurred for a few beads then quickly stopped (Fig. S1). To investigate the cause, we calculated the free energy of two laterally-interacting filaments as a function of their overlap (Fig. S2). We found that the system’s energy minimum decreased much faster than the energy maximum as the number of lateral bonds increased, resulting in an increasing energy barrier. At a high number of lateral bonds, the energy barrier was sufficiently high to prevent thermal fluctuations from breaking avidity and allowing further sliding.

#### Filament depolymerization

In order to maintain a low number of lateral bonds, and therefore a low energy barrier, we implemented filament depolymerization during filament sliding (Fig. S2). Considering that ring integrity would be lost if depolymerization were faster than sliding, we calculated the upper limit of the filament sliding rate. Due to the presence of a large turgor pressure, the membrane constriction rate and therefore the filament sliding rate are limited by the rate of inward cell wall growth. In our simulations, the average inward growth rate of the cell wall was calculated to be *v*_*radial*_ = 14 nm/s, equivalent to a circumference reduction rate of *v*_*cir*_ = *2πv*_*radial*_ = 88 nm/s. As the average length of the filaments was set to be 176 nm (40 monomers), a complete ring of radius 234 nm would be composed of at least 9 filaments. The maximum filament sliding rate was there-fore *v*_*cir*_/9 ∼ 10 nm/s. This value was equivalent to a critical depolymerization rate of *r*_*c*_ = 2.3 beads/s considering that the distance between adjacent beads was *l*_*z*_ = 4.4 nm. As expected, in simulations with a depolymerization rate of 5 beads/s, which was faster than the critical rate *r*_*c*_, rings were quickly broken, limiting constriction (Fig. 2E). Simulations with a depolymerization rate of 1 bead/s resulted in a deep constriction and much slower loss of ring integrity (Fig. 2F).

#### Filament treadmilling

Since FtsZ filaments treadmill around the cell circumference (*14, 15*), we speculated that treadmilling might provide an alternative mechanism to maintain a low degree of lateral contact to sustain sliding. We therefore removed depolymerization and implemented filament treadmilling. Note that the treadmilling direction was randomly chosen for each filament in accordance with the experimental observation of FtsZ treadmilling in both directions in cells (*14, 15*). As expected, simulations with a treadmilling rate of 1 bead/s resulted in constriction (Fig. 2G, H). However, we also observed the formation of bundles of filaments (Fig. S3). Detailed analysis revealed three scenarios for how filaments treadmill with respect to one another (Fig. S4). First, if two filaments in lateral contact treadmill away from each other, their lateral contact is reduced, leading to further filament sliding, similar to the effect of depolymerization. Second, if two filaments in lateral contact treadmill in the same direction, the degree of lateral contact does not change. Third, if two filaments in lateral contact treadmill toward each other, their lateral contact increases, further increasing avidity and gradually forming a bundle as another filament comes in.

### Filament bending model

For simplicity, in this model we assumed that FtsZ filaments bend without twisting (Fig. S5). To prevent twisting, we represented each FtsZ monomer as a cube with vertices connected by springs (Fig. S6). To enable the filament to bend, the two springs on the C-terminal side were replaced with springs of a longer relaxed length while the two on the N-terminal side were replaced with springs of a shorter relaxed length (see **Methods/Filament bending model** for details). Note that in most figures of this model, the cubes are shown as beads for simplicity.

#### GTP-hydrolysis-induced bending

FtsZ filaments are thought to be straight when in the GTP-bound state and switch to a bent conformation upon GTP hydrolysis (*8*). It is likely that both GTP- and GDP-bound states co-exist in real cells, but for simplicity in our simulations, all filaments were initiated in the GTP-state and switched to the GDP-state once the simulation started. To test whether bent filaments can deform the membrane as observed *in vitro* (*35*–*37*), we started with a membrane of diameter 500 nm (Fig. 3A). We implemented a preferred curvature of the filaments 10 times that of the membrane such that bent filaments could form rings 50 nm in diameter, approximately twice the size of FtsZ mini-rings, the smallest rings that have been observed *in vitro* (*28*). Our simulations did not lead to membrane deformation, however. Instead, filaments rolled as they bent, eventually forming arcs on the plane of the membrane (Fig. 3B). This topology is more energetically favorable and is consistent with the prediction by Erickson et al. (*8*). Our results therefore suggest that for filament bending to deform the membrane, the cell must have a mechanism to keep the bending in the division plane.

**Figure 3:**
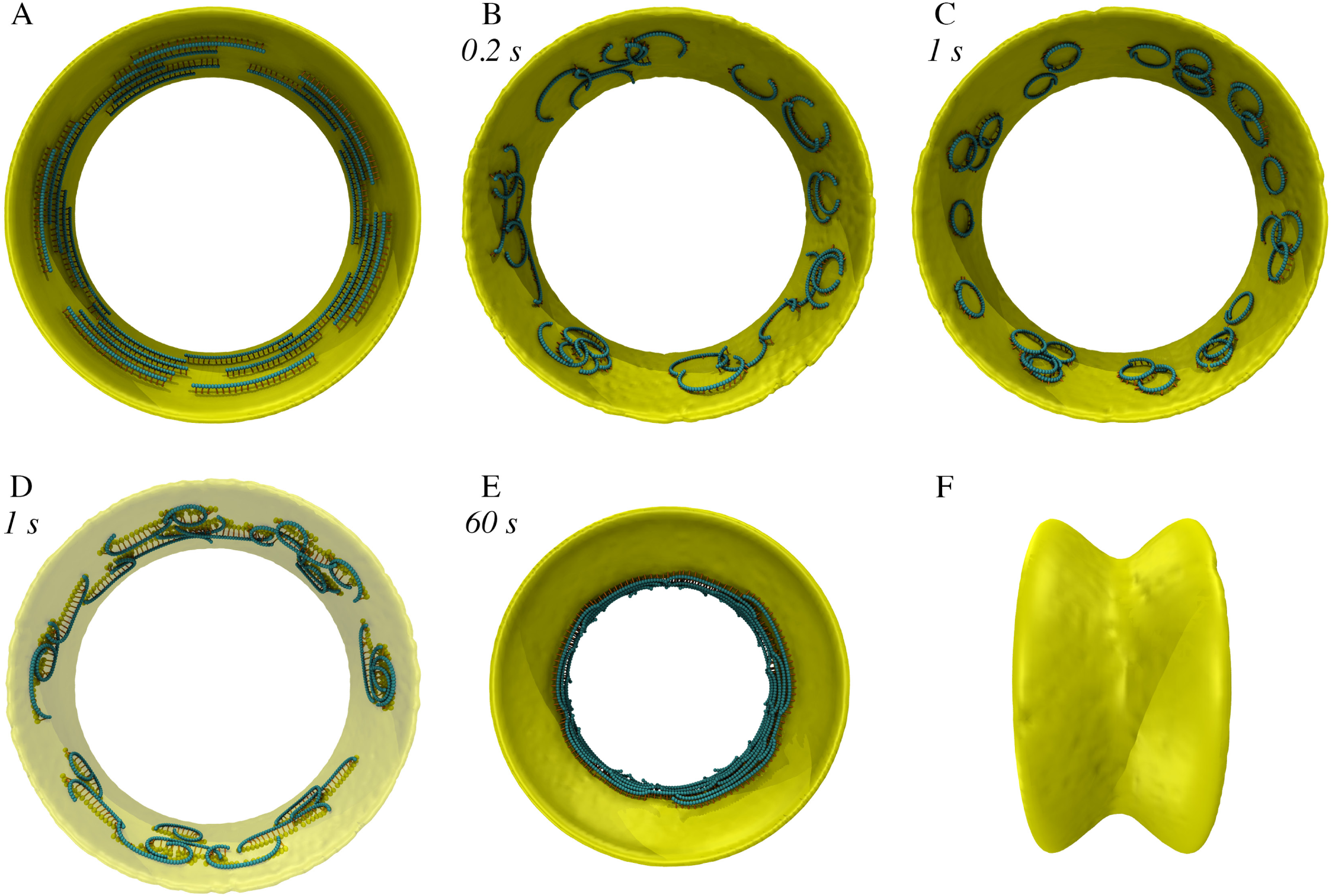
Simulation results of the filament bending model. Italic font indicates simulation times. (A) The initiated system with FtsZ filaments in the GTP-state running circumferentially. (B) After the filaments were switched to the GDP-state, they did not constrict the membrane but instead rolled to bend on the plane of the membrane. (C) Implementing treadmilling did not prevent rolling, but filaments treadmilled in circles on the plane of the membrane. (D) Aligning membrane beads (yellow) connected to the same filament to the circumferential direction did not prevent filament rolling but only stretched the filament circles into a more elliptical shape. Note that, except for beads connected to filaments, the whole membrane (visualized as a surface) is shown with low opacity. (E) The presence of rigid linkers that prevented rolling allowed filaments to exert force on the membrane. Treadmilling resulted in uniform membrane constriction. (F) Side view of (E).

To test whether filament rolling was still energetically favorable with dynamic filaments, we implemented filament treadmilling with a rate of up to 14 beads/s. The resulting simulations showed that treadmilling did not prevent rolling, but rather caused filaments to form dynamic circles (Fig. 3C), similar to experimental observations of reconstituted FtsZ filaments on a flat membrane (*39*).

#### Circumferentially constrained linkers

Searching for mechanisms to prevent filament rolling, we speculated that connecting filaments to the cell wall might reduce rolling since glycan strands could provide a circumferential template in real cells. To test this hypothesis, we constrained membrane beads connected to the same filament to the circumferential direction (see **Methods/Circumferentially constrained linkers** for details). Simulations showed that this circumferential constraint still failed to prevent filament rolling, however (Fig. 3D; Fig. S7A). Because both the filaments and linkers were flexible, partial rolling could still occur, causing the filaments to treadmill in elliptical tracks. In addition, we observed that filaments were pulled close to the membrane (Fig. S7B, C), in contrast to the large distance of ∼16 nm between filaments and the membrane observed in cryo-ET studies of real cells (Fig. S7D) (*9, 10*).

#### Rigid linkers

We reasoned that flexible linkers alone cannot prevent filament rolling and maintain the large distance between the filaments and the membrane. Since many proteins are thought to bind FtsZ *in vivo*, for example FtsA, ZipA, and ZapABCDE (*11*–*13, 40*–*45*), we hypothesized that these proteins can form a rigid complex connecting the filament to the membrane. To maintain the filament-membrane distance, we first modified the linker (originally modeled as two springs joined at a central bead) to be a single spring (Fig. S8) (see **Methods/Rigid linkers** for details). We then constrained this linker to the radial direction of the membrane. We assumed this linker could prevent filament rolling and therefore constrained the bending within the division plane. Simulations of this system showed that the filaments could now pull the membrane down and, due to filament treadmilling, membrane constriction occurred uniformly (Fig. 3E, F).

We initially assumed that FtsZ filaments would bend to a high curvature, consistent with the observation of 24 nm mini-rings (*28*). However, in several studies, FtsZ filaments were observed to form large rings *in vitro* with diameters ranging from 150 – 1000 nm (*30*–*34, 46*–*48*). We therefore ran simulations in which the preferred curvature of the filaments was only twice that of the membrane. Specifically, the membrane was originally 500 nm in diameter and the filaments could form rings 250 nm in diameter. As we expected, these simulations did not result in membrane constriction (Fig. 4A), confirming that a highly bent filament is required for the bending-driven constriction mechanism to work.

**Figure 4:**
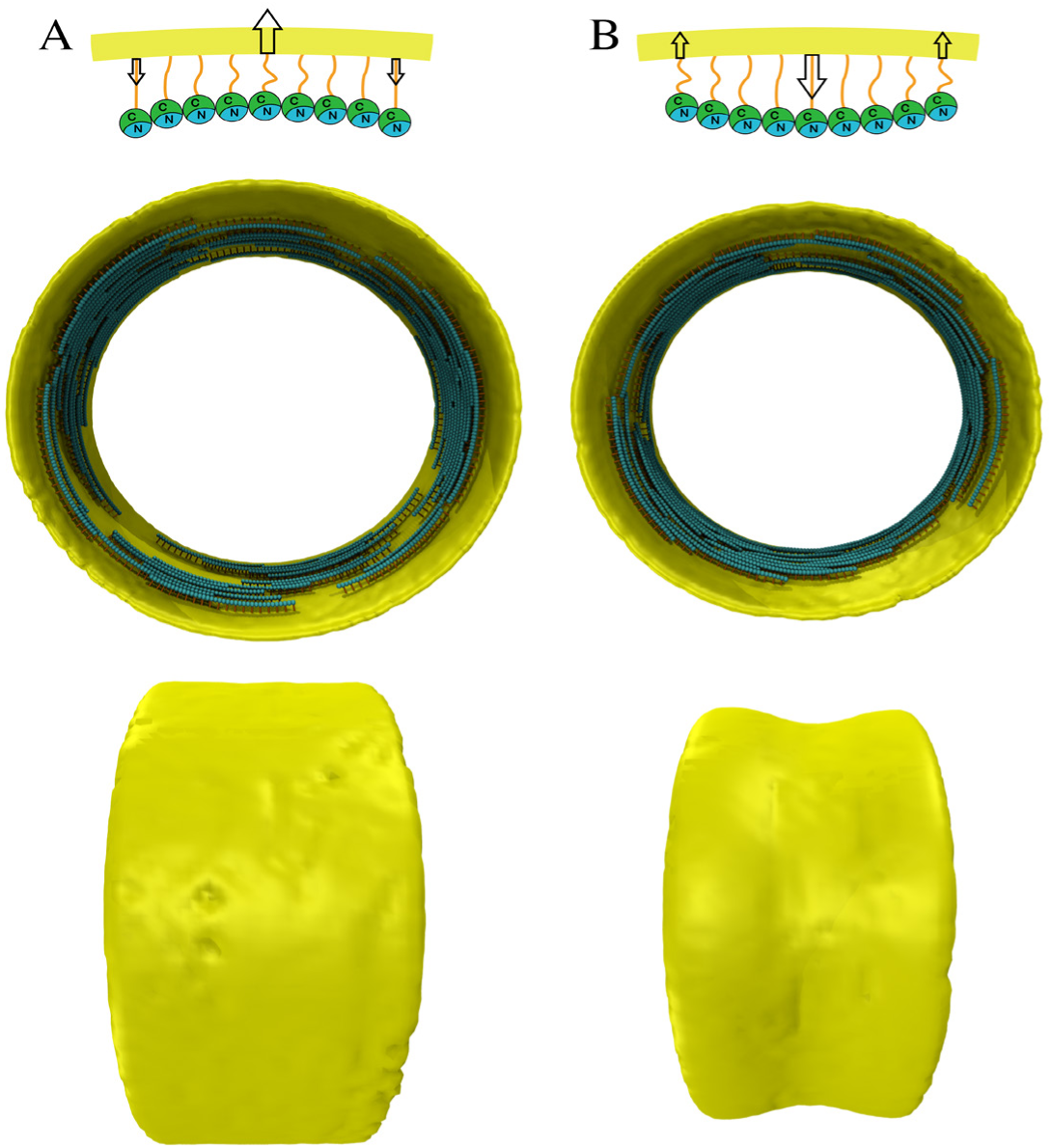
In-sync bending vs reverse bending. (Top) Schematic of filament bending that is in-sync (A) or reverse (B) to membrane bending. In the reverse bending, the C-terminus is on the inner curvature of the filament. Arrows indicate directions of forces exerted on the membrane by the filament. (Middle) Simulation results after 100 sec of systems in which the membrane initially had a diameter of 500 nm and the preferred curvature of the filament was 1/250 nm^−1^ (twice that of the initial membrane). The in-sync bending did not result in constriction, but the reverse bending did. (Bottom) Side views.

#### Reverse bending

Using physical reasoning, Erickson et al. (*8*) pointed out that even mini-rings of 24 nm diameter (*28*) could only constrict the membrane to an ∼50-nm diameter, taking into account the FtsZ-membrane distance of ∼16 nm and assuming FtsZ can fully bend to its preferred curvature. However, except for cases where the filaments were allowed to roll on the membrane, in our simulations we never observed filaments that fully bent to their preferred curvature. Intuitively, the constrictive force depends on the difference in curvature between the membrane and the filament. Therefore, as the membrane becomes smaller, the constrictive force is reduced. When the resistance from the membrane bending stiffness and turgor pressure balances the constrictive force, constriction would stop.

The above reasoning assumes that the C-terminus of FtsZ, which is connected to the membrane via linker proteins, is on the outer curvature of the bent filament. However, Li et al. found evidence that the C-terminus is instead on the inner curvature of the bent filament (*49*). The authors assumed that both the filament and the membrane bend in the same direction and therefore speculated that the C-terminal linker would wrap around the filament to connect with the membrane (Fig. S9). We found this arrangement unintuitive, and instead reasoned that if the C-terminus is on the inner curvature, the filament would bend in the opposite direction of the membrane (Fig. 4B). In this case, even slightly bent filaments might be able to constrict the membrane. To test this hypothesis, we repeated simulations in which the membrane was originally 500 nm in diameter and the filaments could form rings 250 nm in diameter, with the modification that the filaments were now modeled to bend in the opposite direction of the membrane and the C-terminus was now on the inner curvature. As we expected, reverse bending of filaments with even a low curvature resulted in membrane constriction (Fig. 4B).

#### Single-filament constriction

A recent cryo-ET study from our lab showed that in several species, constriction initiated with a single FtsZ filament visible (*27*). Since a single filament can only exert force locally on the membrane, we speculated that the filament must connect to the cell wall. If so, as the filament bent, its end-to-end distance would be reduced, locally relaxing the pressurized cell wall (Fig. S10). This relaxation of the cell wall creates the condition that our previous simulations suggest would allow new cell wall of a smaller diameter to be incorporated (*4*). We therefore ran simulations of the reverse bending model in which there was only a single FtsZ filament with a preferred curvature of 1/32 nm^−1^ and observed membrane constriction (Fig. 5).

**Figure 5:**
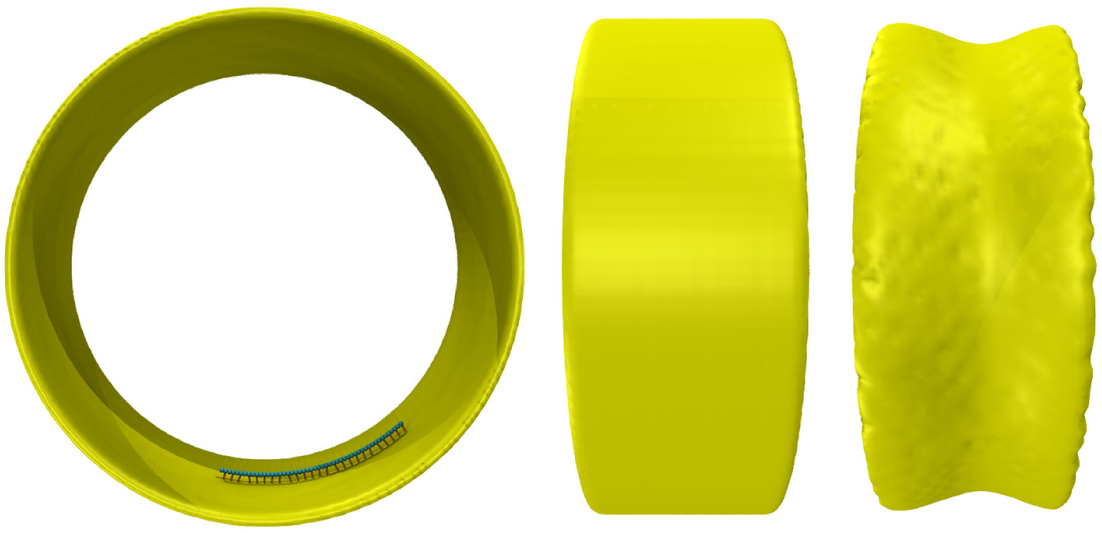
Single-filament constriction. (Left) An axial view of the initiated single-filament system. (Middle) A side view of the initial system. (Right) A side view of the system after 100 sec of the simulation time shows that the single FtsZ filament could constrict the membrane through reverse bending and treadmilling.

## DISCUSSION

How bacterial cells generate a constrictive force for division without motor proteins remains a puzzle. To explore possible mechanisms, here we designed software ZCONSTRICT and performed simulations to test two popular conceptual models.

In the filament sliding model, overlapping FtsZ filaments form lateral bonds. As filaments interact to increase the number of lateral bonds, the ring tightens, constricting the membrane. For this model to work, we found that it requires the following conditions. First, a long-range attractive force between the filaments is needed to first form and then increase the number of lateral bonds between the filaments. While we implemented a Lennard-Jones potential in our simulations, we are not aware of any mechanism that generates such an interaction in real cells. Second, since long-range attraction quickly collapses parallel filaments into a bundle with high avidity between the filaments, a mechanism that separates filaments into multiple rings is needed to allow filament sliding. For simulation purposes, we assumed the action of a protein(s) that separates rings, but we are not aware that such a protein has been identified in cells. Third, as sliding increases lateral contact and avidity, a mechanism such as depolymerization or treadmilling is needed to maintain a low number of lateral bonds to sustain sliding. Previous work by Lan et al. using Monte Carlo simulations predicted that sliding can occur since the lateral energy of the system decreases as the lateral contact increases (*26*). The authors only evaluated the energy of states where beads on one filament are in perfect register with those on the partner filament and their evaluation is consistent with ours for these “in register” states. However, similar to the evaluation by Erickson (*50*), our calculations showed that the energy barrier between adjacent “in register” states (in other words, the cost to break the bonds and transition through an intermediate state where the beads are not in register) increases with the number of lateral bonds, eventually preventing sliding. Finally, sliding can generate ring tension only if filaments overlap to form a complete ring. Cryo-ET imaging studies from our lab and others observed complete rings in mid and late stages of constriction (*10, 27*). However, in early stages in several species, we observed constriction without a complete ring (*9, 27*), suggesting that at least initially a mechanism other than filament sliding is responsible for membrane constriction.

In the filament bending model, FtsZ filaments hydrolyze GTP to switch from a straight to a bent conformation. Due to the curvature mismatch between the filament and the membrane, bent filaments exert a constrictive force on the membrane. For this model to work, we found that it requires the following conditions. First, a mechanism that constrains the bending to the division plane is needed to prevent the filaments from rolling to bend in the plane of the membrane (since bending without exerting force on the membrane is more energetically favorable). This was predicted by Erickson et el. (*8*). While we assumed that a rigid linker formed by several FtsZ-binding proteins prevents filament rolling, we are not aware of any evidence for such a linker *in vivo*. Second, a large difference in curvature between the membrane and the filaments is needed for filament bending to overcome resistance from membrane stiffness and turgor pressure. If filaments bend in the same direction as the membrane, we found that a small preferred curvature of FtsZ was not adequate to initiate membrane constriction. Even if the preferred curvature were sufficiently large to start constricting the membrane, constriction would stop once the membrane curvature approached the preferred curvature of FtsZ. In contrast, our simulations showed that constriction can start and sustain with a small FtsZ curvature if filaments bend in the direction opposite that of the membrane. This could help explain the observation that the C-terminus of FtsZ, which is attached via linkers to the membrane, is on the inner curvature of the bent filament (*49*). A more recent study, however, showed evidence that the C-terminus is on the outer curvature of the bent filament (*51*),so evidence for both in-sync bending and opposite bending exists (*49, 51*).

The filament bending model could account for observations of initial constriction with a single filament (*27*). Note that FtsZ filaments are highly dynamic, so it is possible that other filaments depolymerized just before the cryo-ET snapshots were captured, but the observation strongly suggests that complete rings are not required. In our simulations here we speculated that for incomplete rings to be able to constrict the membrane, the filaments must be connected to the cell wall so that their bending can overcome the effect of turgor pressure and locally relax glycan strands, a condition that our previous simulation work predicted to be required for making hoops of new cell wall that are smaller than existing hoops (*4*). While this FtsZ-cell wall connection assumption provides a possible mechanistic explanation for the single-filament constriction observed experimentally, we are not aware of clear evidence for such a connection in real cells. In later stages of constriction, more filaments and complete rings are present (*10, 27*), so it is also possible that a combination of mechanisms contributes to robust constriction *in vivo*.

While it is widely thought that GTP hydrolysis drives filament bending, the size mismatch between FtsA and FtsZ has also been speculated to cause filament bending (*52*). In this model, FtsA forms a filament parallel to the FtsZ filament. The difference in the FtsA monomer length (4.8 nm) and the FtsZ monomer length (4.4 nm) and the one-to-one connection between the two causes bending. We estimate that a tight connection between the two filaments would force the FtsZ filament to form a circle ∼50 nm in radius. Since the distance between the membrane and FtsZ is ∼16 nm (*9, 10*), this circle could potentially constrict the membrane to a radius of ∼66 nm. Note that we are not aware of any evidence supporting or refuting the possibility of copolymerization of FtsA and FtsZ in real cells.

Likewise, FtsZ might generate a constrictive force by a completely different mechanism. For example, FtsZ has been recently shown to treadmill around the cell circumference (*14, 15*). While treadmilling was speculated to be a mechanism to uniformly distribute new cell wall material around the division site, such dynamics have the potential to generate movement of other proteins and consequently a mechanism to generate a constrictive force. Finally, we cannot rule out other force generation mechanisms that do not involve FtsZ, for instance excess membrane synthesis, which has recently been proposed (*53*).

As with any simulation of a conceptual model, our work has limitations. First, we were only able to model a limited number of cellular components: the membrane, cell wall, FtsZ filaments, and linker proteins. Second, we assumed the cell wall does not actively push on the membrane, but simply passively grows in after the membrane. While our previous work provides support for this assumption (*4*), it has not yet been validated experimentally. Third, due to the large scope of our simulations, we could only explore a small parameter space for each conceptual model. While we always subjected our findings to physical reasoning, we cannot rule out the possibility that the findings were specific to the parameter set we tested. Even with these limitations, we hope that our work serves as another example of the power of quantitative 3D modeling to test conceptual models and provide insights into basic principles that remain experimentally inaccessible.

## METHODS

In this section, we describe the design of the software ZCONSTRICT and implementation of assumptions of our simulations.

### Membrane

Similar to our previous work (*5*), we modeled the membrane as a sheet of beads originally forming a cylinder (Fig. 1). In all of our simulations, the original membrane cylinder was 160 nm wide and 250 nm in radius. The bead size was set at *d*_*mb*_ = 8 nm such that if the distance *d* between two beads was smaller than *d*_*mb*_, a force *F*_*push*_ = *k*_*pair*_ *(d*_*mb*_ *– d)*^*2*^ was applied on each bead to push them apart, where *k*_*pair*_ = 1 pN/nm^2^ was the pair-wise force constant. To preserve membrane integrity, a pair-wise attraction was implemented between neighboring beads such that if the distance *d* between two beads was larger than *d*_*pair*_ = 16 nm, a force *F*_*pul*_ = *k* _*pair*_ *(d – d* _*pair*_ )^*2*^ was applied on each bead to pull them together. To implement membrane fluidity, the pair list was recalculated every 10^4^ time steps. As different pairs were formed based on the updated positions of the beads, the beads were allowed to move along the membrane. In our previous membrane model (*5*), a pair was removed from the pair list if it was crossed (looking outward from the center) by a shorter pair. In the current model, to avoid a large change in force on the system as the pair list was recalculated, one degree of crossing was allowed, meaning that a pair was removed from the pair list only if it was crossed by more than one shorter pair.

To implement membrane bending stiffness, if four beads were part of five pairs (Fig. S11), they were constrained to the same plane. If the two diagonals were separated at a distance *d*, a force *F*_*mb*_ = *k*_*mb*_*d* was exerted on the beads to pull the two diagonals together, where *k*_*mb*_ = 8 pN/nm was a force constant. Similar to our previous membrane model (*5*), to prevent boundary artifacts, we implemented a periodic boundary condition such that images of the beads on one longitudinal edge were translated to interact with the beads on the other edge. The translational distance was the same as the membrane width.

### Filament sliding model

In this model, the FtsZ filament was modeled as a chain of beads, each bead representing one FtsZ monomer. The adjacent beads were connected by springs of a relaxed length of *l*_z_ = 4.4 nm (*28*) and a spring constant chosen to be *k*_z_ = 0.5 nN/nm, which was sufficiently large to prevent significant stretching or compression of the filament. If the filament bent an angle *θ* at a bead, an energy *E*_*θ*_ = *k*_*θ*_ (*θ* – *θ*_*0*_)^2^/2 was added to the system, where *θ*_*0*_ = 0° was the relaxed angle. We derived the bending stiffness constant as *k*_*θ*_ = *k*_*B*_ *TL* _*p*_ */ l*_*z*_ = 3.8×10^−18^ J, where *k*_*B*_ is the Boltzmann constant,*T* = 295 K is the room temperature, and *L*_*p*_ = 4 μm is the filament persistence length (*54*). Note that much smaller values of FtsZ persistence length ∼100 nm have also been reported (*18, 33*).

We implemented a Lennard-Jones potential as a long-range interaction between beads on different filaments (note that interaction of beads on the same filament was excluded). Specifically, a force between two beads at a distance *d* was calculated as:

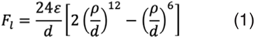

where the distance at which the potential is zero was set to be *ρ* = 6.5 nm, ∼ the distance between paired filaments observed via cryo-ET of dividing cells (*10*). Varying the depth of the potential *ε* in the range from 10^−19^ to 10^−18^ J with a step size of 10^−19^ J, we found that the lower bound was too weak to result in filament sliding while values larger than 5×10^−19^ J resulted in filaments twisting around each other. We therefore chose *ε* = 5×10^−19^ J, the largest value that induced filament sliding without causing filament twisting.

To implement filament depolymerization and treadmilling, the FtsZ filament was modeled to be polar with one end tracked to be “plus” and the other “minus.” Depolymerization was modeled by removing a bead at the minus end with a rate varying from 1–10 beads/s. Treadmilling was modeled by adding a bead to the plus end and removing a bead from the minus end at the same rate, which was varied from 1–14 beads/s.

### Filament bending model

We assumed that the filament bends in the same plane without twisting. To implement this assumption, we modeled the filament as a chain of cubes, with each cube representing one FtsZ monomer (Fig. S6). For convenience, the beads on each cube were denoted 1, 2, 3, 4, 5, 6, 7, 8. Note that the beads on the interface of two adjacent cubes were shared, meaning beads 5, 6, 7, 8 of one cube were the same as beads 1, 2, 3, 4 of the other. The C-terminal face contained beads 1, 2, 6, 5 and the N-terminal face contained 4, 3, 7, 8. To maintain the width of the filament, the beads on the two cross-sectional faces (perpendicular to the long axis; one containing 1, 2, 3, 4 and the other 5, 6, 7, 8) were connected by springs of relaxed length *l*_*z*_ = 4.4 nm and spring constant *k*_*z*_ = 0.5 nN/nm (Fig. S6A). Note that calculation of the spring force on the interface of two adjacent cubes was done only once for each spring. To maintain the length of the filament, the two cross-sectional faces were connected by two springs of relaxed length *l*_*z*_ and spring constant *k*_*z*_. One spring connected edge 1-4 to edge 5-8 and the other connected edge 2-3 to edge 6-7 (Fig. S6B).

To model filament bending, the beads on the C-terminal face were connected by two springs of relaxed length *l*_*C*_ and spring constant *k*_*b*_, one spring connecting bead 1 to bead 5 and the other connecting 2 to 6 (Fig. S6B). Likewise, on the N-terminal face, one spring of relaxed length *l*_*N*_ and spring constant *k*_*b*_ connected 3 to 7 and another (identical) spring connected 4 to 8. When the filament was in a straight conformation (supposedly GTP-bound), both *l*_*C*_ and *l*_*N*_ were set to be *l*_*0*_, which was the distance between the two centers of the two cross-sectional faces (Fig. S6B). Note that *l*_*0*_ was not constrained to be constant but varied with the stretching/compression of the filament. When the filament was in a bent conformation (having already hydrolyzed GTP) with a preferred angle *θ*_*β*_ between adjacent monomers, *l*_*Ψ*_ was set to be *l*_*0*_ + *Δl*_*0*_ and *l*_*N*_ was set to be *l* - *Δl* where 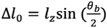 (Fig. S6C). In the case that reverse bending was assumed, *l*_*C*_ was set to be *l*_*0*_ – *Δl*_*0*_ and *l*_*N*_ was set to be *l*_*0*_ + *Δl*_*0*_. In our simulations, we varied *θ*_*b*_ between 1° and 20°. Considering the distance between adjacent subunits *l*_*z*_ = 4.4 nm, bent filaments that form circles with diameters of 250 and 50 nm correspond to *θ*_*b*_ = 2° and 10°, respectively. The spring constant *k*_*b*_ was calculated by setting the energy stored in four springs as their length was deformed an amount *Δl*_*0*_ to be the bending energy of the filament with bending stiffness *k*_*θ*_ (the same constant as in the filament sliding model) as follows:

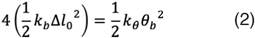

Note that *θ*_*b*_ was converted to radians when the bending energy was calculated.

To prevent filament twisting, the two diagonals of each face were constrained to the same length such that if their lengths differed an amount *Δl*, a force *F*_*diag*_ = *k*_*z*_*Δl* was applied on the four beads to restore the lengths to the same value (Fig. S6D).

### Filament-membrane connection

Each filament was connected to the membrane via linkers (Fig. 1). Each linker was composed of two identical springs of a spring constant of *k*_*lk*_ = 20 pN/nm and a relaxed length of *l*_*lk*_ = 8 nm, making the distance from the filament to the membrane 16 nm as reported experimentally (*9, 10*). One spring was connected to an FtsZ monomer and the other to a membrane bead and they were joined by a bead representing FtsA or ZipA. In the case of the filament bending model, because each FtsZ monomer was represented as a cube, the associated linker was linked to the cube’s center.

### Circumferentially constrained linkers

To constrain the linkers connected to the same filament to the circumferential direction, if two membrane beads that were connected to two adjacent linkers were separated at a distance *d* along the axis perpendicular to the circumferential direction (Fig. S12), a restoring force, *F*_*c*_ = –*k*_*c*_*d*, where *k*_*c*_ = 20 pN/nm was a force constant, was applied to align them back to the circumferential direction.

### Rigid linkers

To make the linker rigid, the original linker model as two springs of a relaxed length of 8nm and a spring constant *k*_*lk*_ = 20 pN/nm joined at a central bead was replaced with a single spring of 16 nm long and the same spring constant *k*_*lk*_. To constrain the linker to the radial direction of the membrane, if the two ends were separated at a distance *d* along the axis that was perpendicular to the preferred (radial) direction (Fig. S12), a restoring force, *F*_*r*_ = –*k*_*r*_*d*, where *k*_*r*_ = 3 pN/nm was a force constant, was applied on the ends to align them back to the radial direction. To constrain filament rolling, the springs connecting the C- and N-terminal faces of the FtsZ-monomer cubes, specifically those that connect bead 1-4, 2-3, 5-8, and 6-7 (Fig. S6), were constrained parallel to the division plane (the plane perpendicular to the cell long axis). If a constrained spring deviated from the division plane, a restoring force of magnitude *F*_*dp*_ = *k*_*dp*_*d*, where *k*_*dp*_ = 20 pN/nm was a force constant and *d* was the projection of the spring length on the cell long axis, was applied on each of the two end beads of the spring to align it to the division plane (Fig. S13).

### Cell wall

Previously, we modeled the cell wall of Gram-negative bacteria as hoops of glycan strands connected by peptide crosslinks in which each strand was modeled as a chain of beads and each hoop was composed of several strands (*3, 4*). In the current work, we did not focus on the dynamics of the cell wall. We therefore simplified our cell wall model to reduce the computational cost. Specifically, the cell wall was modeled as a grid of beads composed of 11 hoops of radius *r*_*g*_ = 265 nm, separated at a distance of 16 nm and originally forming a cylinder 160 nm wide (Fig. 1). There were 104 beads per hoop separated at a distance of 16 nm from each other and 15 nm from the membrane. The beads on the grid were connected to each other by two different types of springs. The glycan springs with spring constant *k*_*g*_ = 100 pN/nm and dynamic relaxed length *l*_*g*_ ran along the circumference. The peptide springs of spring constant *k*_*p*_ = 10 pN/nm and relaxed length *l*_*p*_ = 16 nm ran along the cell’s long axis. Initially, *l*_*g*_ was chosen such that turgor pressure stretched the glycan spring to a length of *l*_*ext*_ = 16 nm. As turgor pressure was chosen to be *P*_*tg*_ = 1 atm, the original *l*_*g*_ was calculated to be

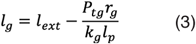

### Turgor pressure

To calculate the force from turgor pressure on the cell wall beads, we first calculated the volume *V* enclosed by the cell wall. To do this, each tetragon was divided into two triangles by one diagonal. Each triangle together with the center point of the cell wall formed a tetrahedron. The total volume *V* was the sum of *V*_*i*_, which was the volume of tetrahedron *i*. The force on each bead *j* was then calculated as *F*_*j*_ = *P*_*tg*_*Σ∇*_*j*_*V*_*i*_. As mentioned above, turgor pressure was chosen to be *P*_*tg*_ = 1 atm.

### Cell wall-membrane connection

We assumed that the cell wall bears all the force from turgor pressure and that this force is transmitted through the membrane and a buffer layer of periplasmic proteins between the membrane and the wall. Similar to our previous model of the membrane (*5*), we modeled the membrane as squeezable, allowing the distance between the membrane and the cell wall to vary around the equilibrium distance of *d*_*w–m*_ = 15 nm. As the distance from a membrane bead to the cell wall deviated *Δd* from *d*_*w–m*_, a force *F*_*w*_* = k*_*w*_ *Δd*^*2*^ (representing the net force from the cell wall and turgor pressure) was applied on the membrane bead to restore it to the equilibrium distance. The force constant was chosen to be *k* _*w*_ = 1.6 pN/nm^2^.

### Cell wall growth

To model cell wall growth, as the membrane was pulled down, the relaxed lengths of two glycan springs connected to the cell wall bead that was closest to the membrane bead were shortened so the wall could move inward to fill the gap. Specifically, every 10^4^ time steps, we calculated the simulated duration *Δt* and the maximal allowed inward displacement of the cell wall *v Δt*, where *v* = 100 nm/s was the limit of the cell wall inward growth rate. If the distance between a membrane bead and the local cell wall surface became larger than *d* _*w–m*_ *+ Δd*_*m*_ where *Δd*_*m*_ = 0.5 nm, the inward displacement of the local cell wall bead *Δd*_*w*_ was chosen as the smaller of either 0.05 nm or *v*_*w*_*Δt*. Note that this rule resulted in an average inward growth rate of the cell wall *v*_*radial*_ = 14 nm/s. The relaxed lengths of the two glycan springs connected to the local cell wall bead were then shortened an amount *πΔd*_*w*_*/*_*b*_, where *N*_*b*_ = 104 was the number of wall beads per hoop.

### Diffusion

As in our previous simulation work (*3*–*5*), we modeled thermal motion of the system by implementing random forces on the beads. We used the Box-Muller transformation to generate a set of Gaussian-distributed random numbers. Specifically, two random numbers of a Gaussian distribution were generated using two random numbers from a uniform 0 – 1 distribution, *u* and *u*, as follows

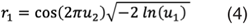

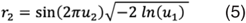

To reduce the computational cost, we did not integrate the Gaussian distribution with a time step to generate random forces. Instead, a force was simply calculated as the product of a random number of the Gaussian distribution with a force constant *k*_*r*_ varying from 1–10 pN.

### Volume exclusion

We implemented a volume exclusion effect between FtsZ beads and membrane beads and between beads on different FtsZ filaments. If the distance *d* from an FtsZ bead to a membrane bead was smaller than *d*_*mb*_ = 8 nm, a force *F* = *k*_*m–z*_*(d*_*mb*_ *– d)*, where *k*_*m–z*_ = 200 pN/nm was a force constant, was applied on the beads to push them apart. In the filament sliding model, the Lennard-Jones potential provided a volume exclusion effect. In the filament bending model, if the distance *d* between the centers of two cubes (of different two filaments) was smaller than *d*_*zz*_ = 6.5 nm, a force *F* = *k*_*zz*_*(d*_*zz*_ *– d)*, where *k*_*zz*_ = 1 nN/nm was a force constant, was exerted on each bead of the two cubes to push them apart.

### System dynamics

As described in detail previously (*3*–*5*), we used the Langevin equation to calculate the system dynamics. Assuming the inertia of each bead was negligible, the displacement of bead *i* at a time step was calculated as

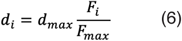

where *F*_*i*_ and *F*_*max*_ were the force on bead *i* and the maximal force on the beads, respectively. To prevent the system from becoming unstable, the maximal displacement of any bead in one time step was constrained to be *d*_*max*_ = 0.01 nm.

The software ZCONSTRICT was written using Fortran (the source codes can be downloaded at https://github.com/nguyenthlam/Zconstrict). The trajectories of the system dynamics were visualized using VMD (Visual Molecular Dynamics) software

## ACKNOWLEDGMENTS

We thank Debnath Ghosal and Andrew Jewett for their helpful discussions and Jane Ding for help setting up simulations on clusters. This work was supported by the NIH (grant R35 GM122588 to GJJ).

