## Supplementary Information for "Simulations of proposed mechanisms of FtsZ-driven cell constriction"

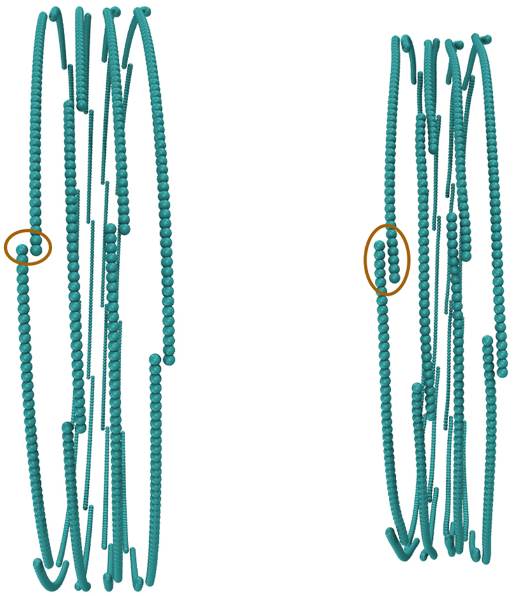

**Figure S1**: Filament sliding model with separated rings. Filaments in each ring were initiated with one bead overlapping (left). Filament sliding occurred but quickly stopped as overlapping reached ~5–6 beads (right). Ellipses indicate examples of overlap.

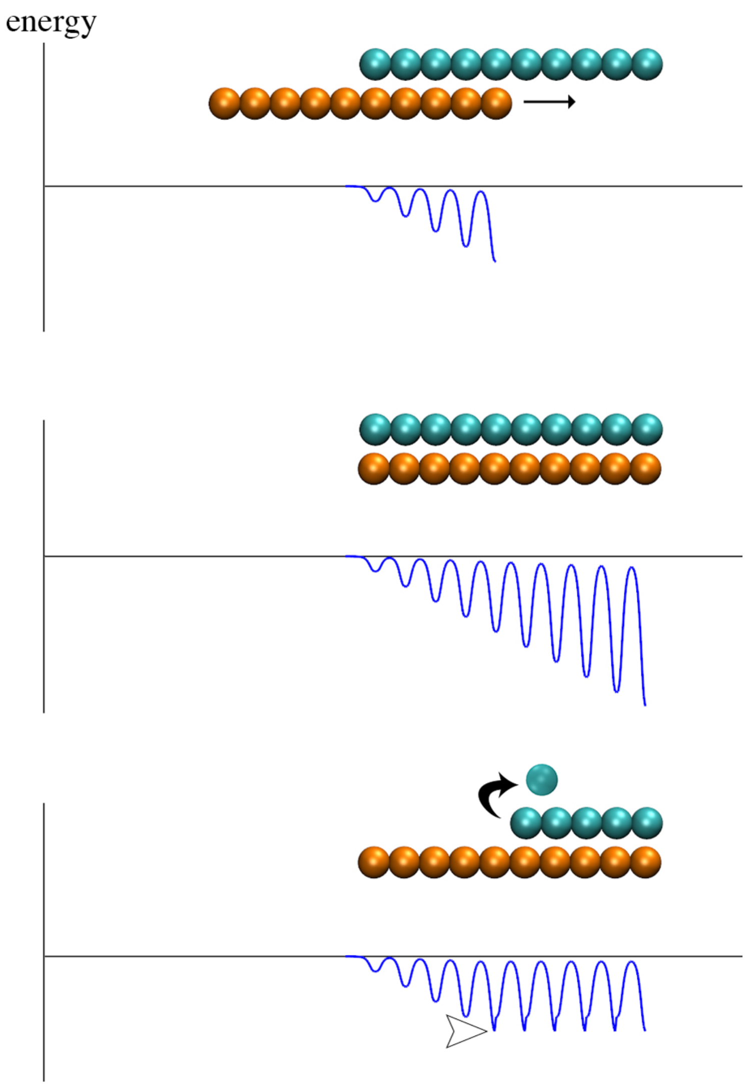


**Figure S2**: Lateral interaction energy of overlapping FtsZ filaments. (Top) As the orange filament slides (indicated by the arrow) with respect to the stationary cyan filament, their interaction energy oscillates with the minimum changing much faster than the maximum, resulting in an increasing energy barrier. (Middle) The energy barrier continues to increase with further filament overlap. (Bottom) If the stationary filament starts to depolymerize (arrow), the energy minimum remains constant (arrowhead), therefore maintaining a low energy barrier as filament sliding continues.

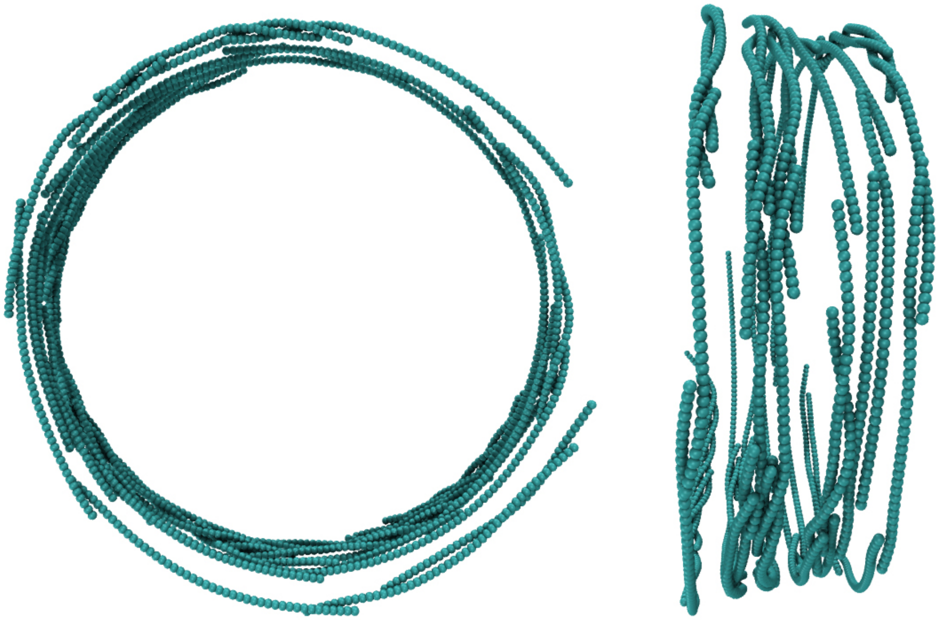

**Figure S3**: Implementation of treadmilling in the filament sliding model. An axial view (left) and a side view (right) of the constricted rings show that several filaments of the same ring overlapped to form bundles.

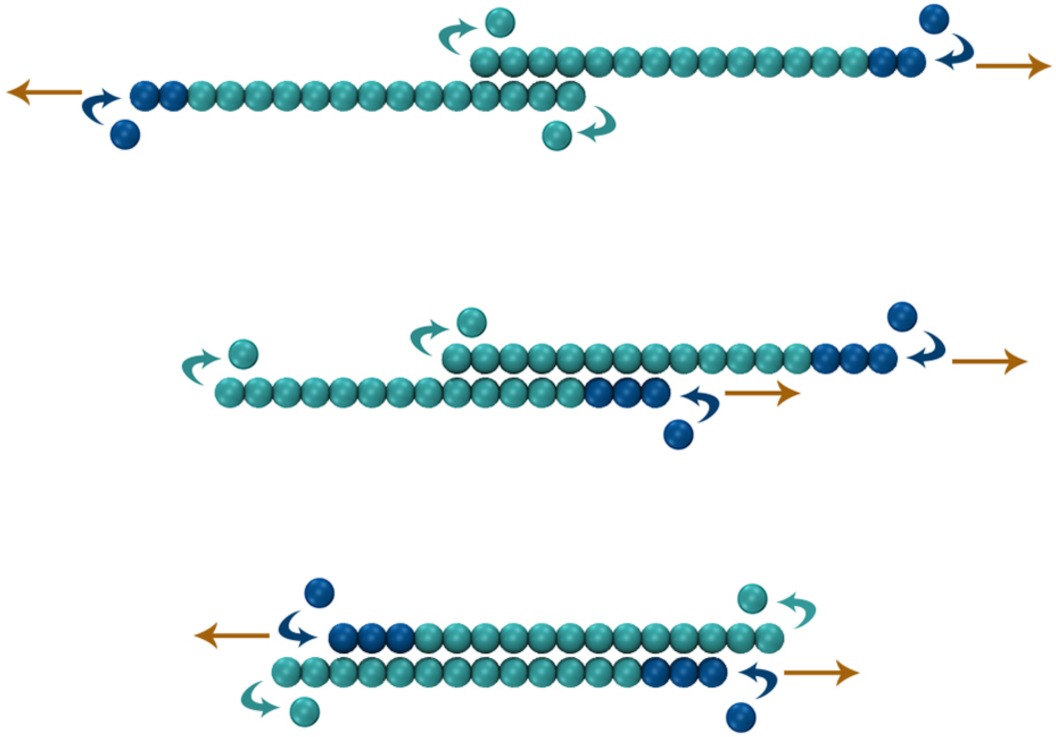

**Figure S4**: Three scenarios of filament treadmilling. Orange arrows indicate treadmilling direction. Dark blue arrows indicate the growing end of the filament. Cyan arrows indicate the shortening end of the filament. (Top) If two filaments treadmill away from each other, the number of their lateral bonds is reduced, allowing further filament sliding (similar to the effect of filament depolymerization). (Middle) If two filaments treadmill in the same direction, their overlap remains constant, causing no effect on filament sliding. (Bottom) If two filaments treadmill toward each other, their overlap increases, further increasing avidity.

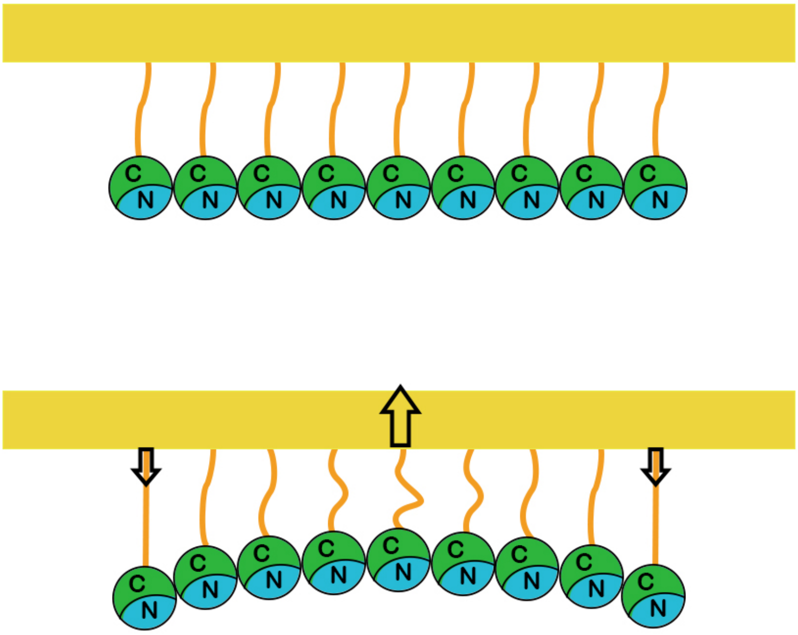


**Figure S5**: Schematic of the filament bending model. FtsZ is connected to the membrane at the C-terminus via flexible linkers. As the filament switches from straight (top) to bent (bottom) conformation, it exerts forces on the membrane. Arrows indicate the force directions.

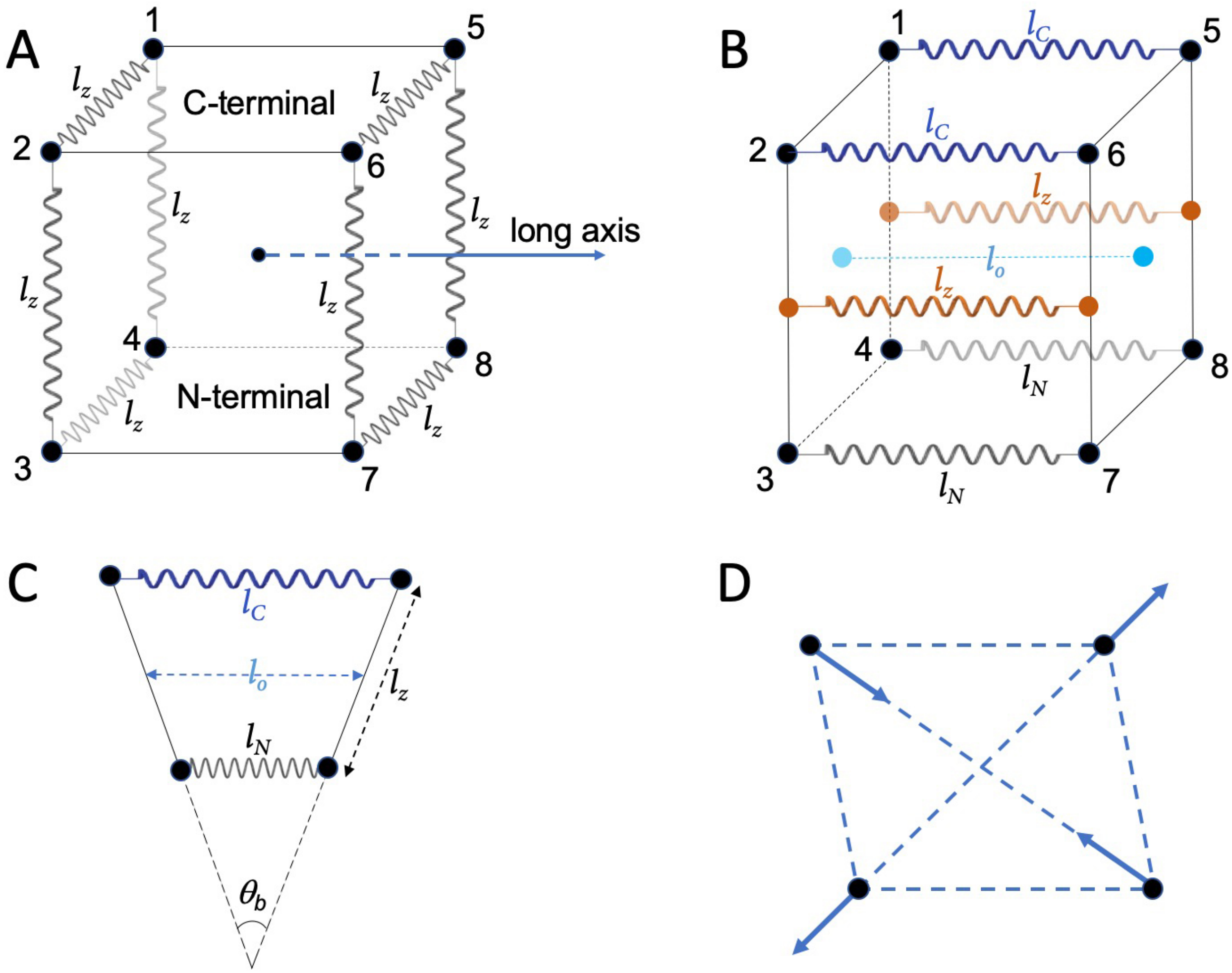


**Figure S6**: Cube model of the FtsZ filament. Each monomer is represented as a cube. (A) The beads on each of the two cross-sectional faces are connected by identical spring of relaxed length $l_z$ = 4.4 nm. (B) The two cross-sectional faces are connected by two springs (orange) of relaxed length $l_z$. One spring connects the center of edge 1-4 to the center of edge 5-8. The other spring connects the center of edge 2-3 to the center of edge 6-7. As the filament is in a straight conformation, the corners are connected by four springs of the same relaxed length, such that both $l_C$ (blue) and $l_N$ (black) are equal to $l_0$, which is the distance between the two centers of the two cross-sectional faces (cyan). (C) To model a bent conformation, $l_C$ and $l_N$ are increased and decreased, respectively. (D) Schematic of forces (arrows) exerted on the beads to restore a distorted face to a square shape.

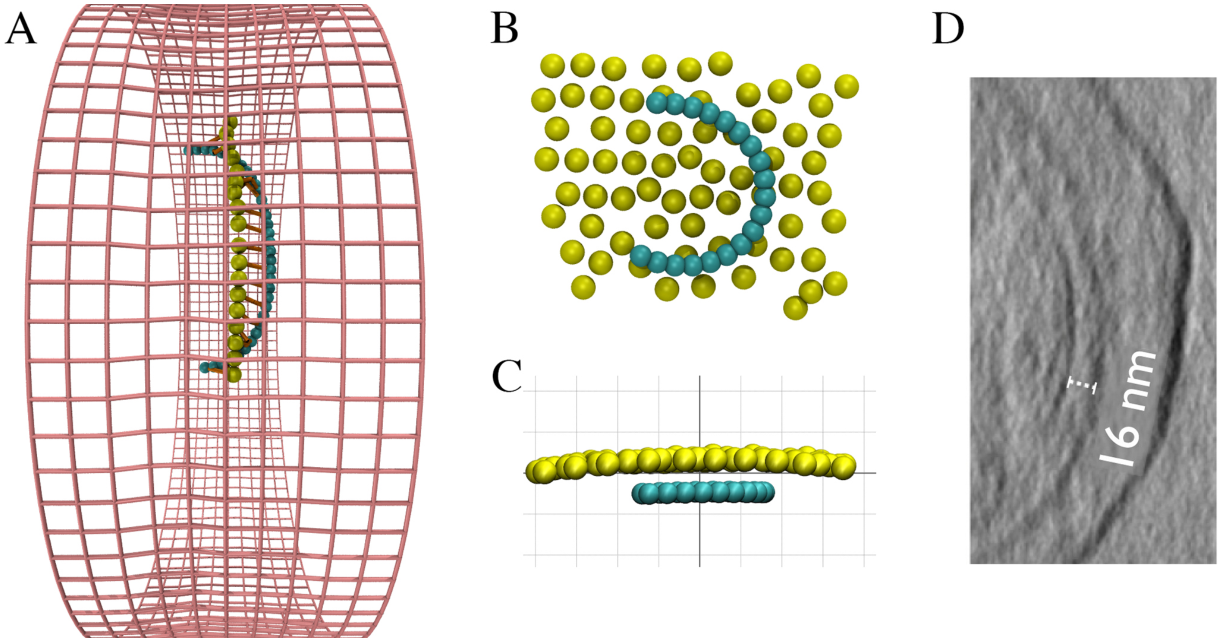


**Figure S7**: Adding a circumferential constraint on the linkers. (A) A filament and its connected membrane beads are visualized together with the cell wall to show that even when membrane beads were aligned to the circumferential direction, the flexibility of the filament and the linkers allowed filament rolling. (B) Front view and (C) side view of a filament with its neighboring membrane beads showing how the filament was pulled close (~8 nm) to the membrane. The grid size in (C) is 10 nm. (D) Adapted from Szwedziak et al. 2014, Figure 1C. A slice through an electron cryotomogram of a dividing cell. The distance from FtsZ to the membrane was measured to be 16 nm, consistent with Li et al. 2007.

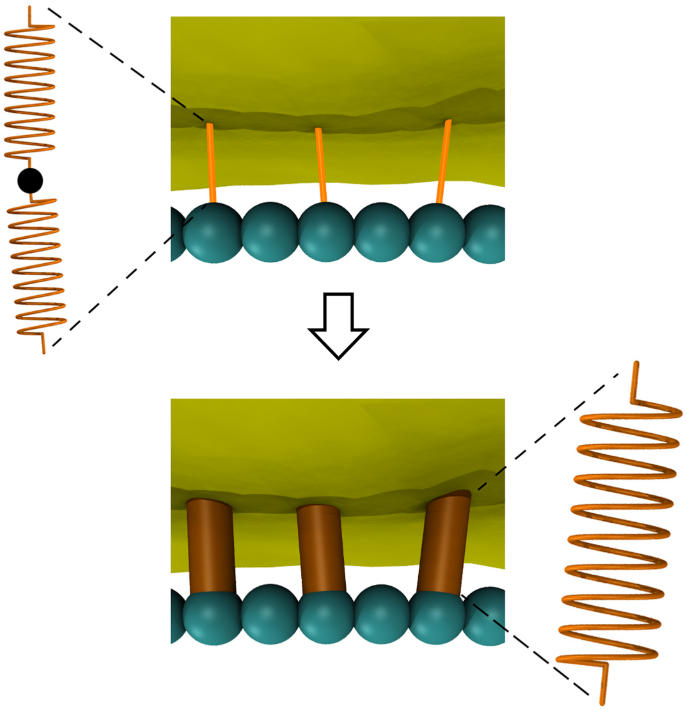

**Figure S8**: Schematic showing replacement of the flexible linker with a rigid linker to maintain the FtsZ-membrane distance and prevent filament rolling.

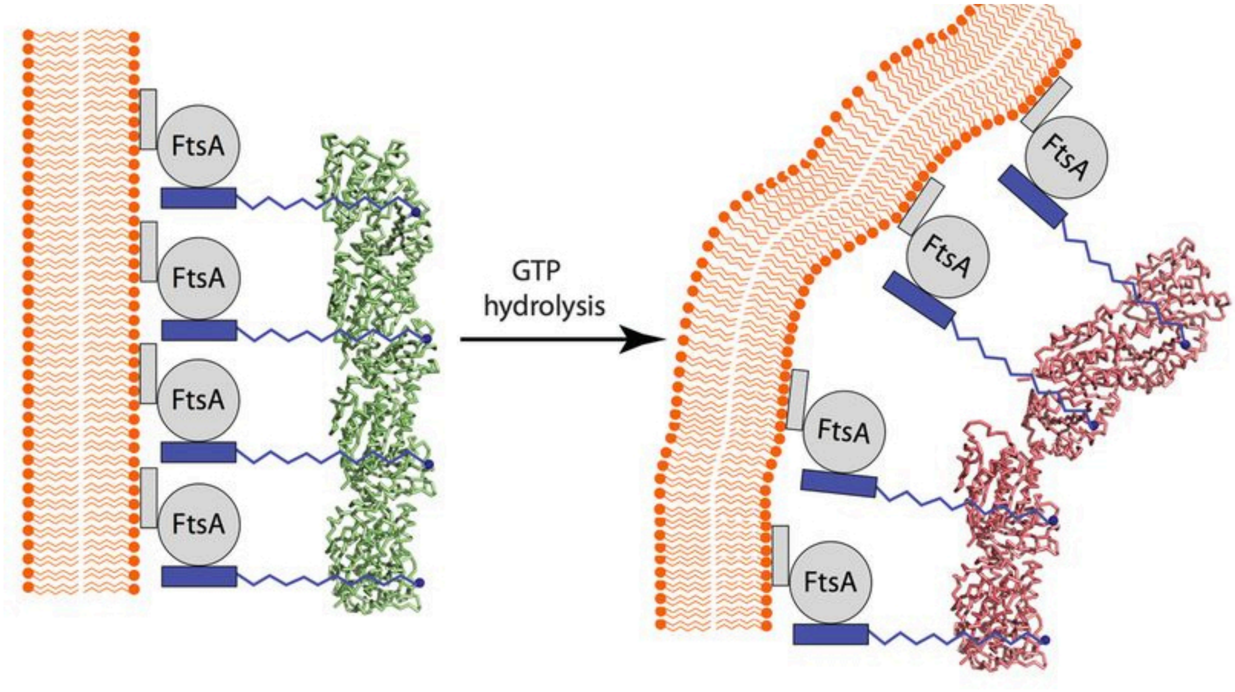


**Figure S9**: Adapted from Li et al. 2013, Figure 3A. The authors show evidence that the C-terminus of FtsZ is on the inner curvature of the bent filament. Assuming the filament bends in the same direction as the membrane, the authors interpreted this to mean that the flexible linker (blue) that connects FtsZ to FtsA must wrap around the filament.

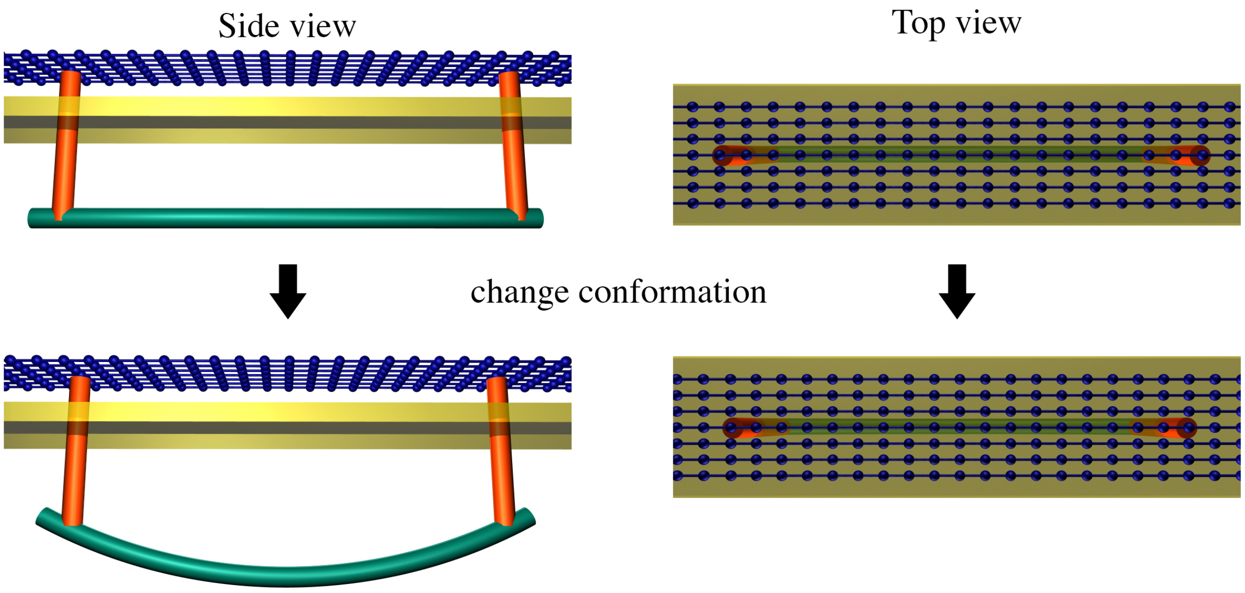


**Figure S10:** Schematic of a connection (orange) between the FtsZ filament (green) and the cell wall (blue). A rigid connection would allow filament bending to squeeze the local cell wall. The membrane is shown in yellow.

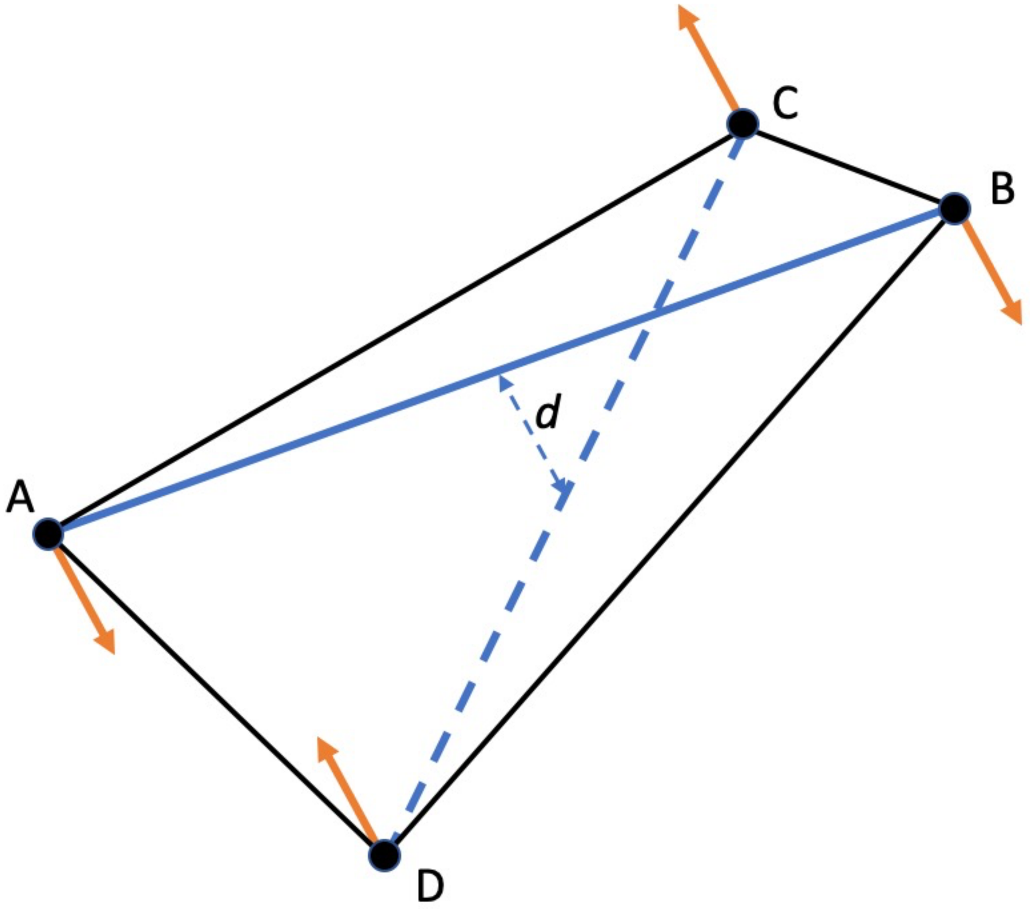


**Figure S11**: Schematic of constraining five pairs (solid lines) formed by four beads A, B, C, D to the same plane. If A is paired with B, both A and B are paired with C, and both A and B are paired with D, then these four beads are confined to the same plane. If line A-B is at a distance *d* from line C-D, a force (orange) is exerted on each bead to pull the two lines into the same plane.

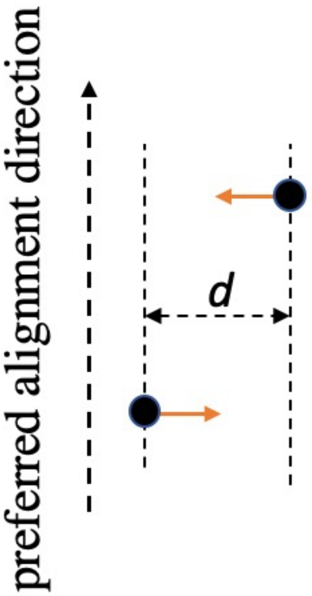


**Figure S12**: Schematic of restoring forces (orange arrows) exerted on two beads to align them to the preferred direction.

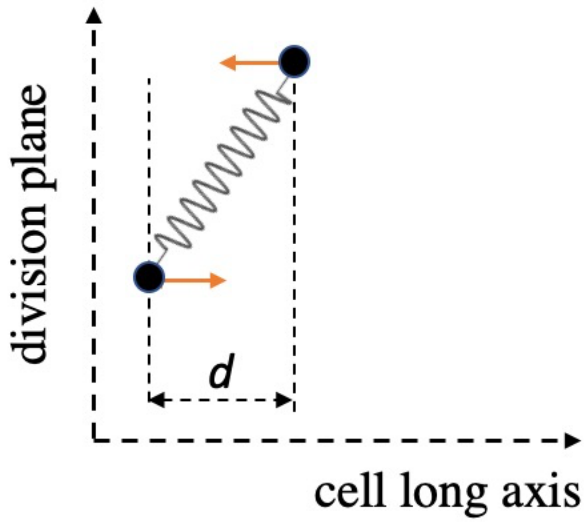


**Figure S13**: If a constrained spring deviated from the division plane, two forces (orange) were exerted on the two end beads to restore its preferred orientation.
